# Combination of indomethacin and temperature produces reliable strobilation in *Cassiopea xamachana* but batch effects create high variability in ephyra outcomes

**DOI:** 10.64898/2026.07.29.741576

**Authors:** Kaden M. Muffett, Jessie G. Puckett, Jacob P. Hébert, Giulio Martino, Maria Pia Miglietta

## Abstract

Indomethacin and temperature shocks are both widely used to induce ephyra production in *Cassiopea* polyps, yet the combined effect of these stimuli on strobilation rate, ephyra viability, and polyp survivorship has not been systematically examined. We designed a full factorial experiment crossing five indomethacin concentrations (0, 10, 25, 50, and 75 µM) with four temperature treatments (22, 25, 28, and 31°C) across three independent experimental trials, exposing a total of 288 individual *Cassiopea xamachana* polyps for 29 days. Overall, 207 of 288 polyps (71.9%) released at least one ephyra. Temperature was the stronger predictor of strobilation: higher temperatures substantially increased the probability of producing a healthy ephyra (log-odds: +0.327 per °C; χ² p = 1.9 × 10^−8^), while indomethacin had a smaller but significant positive effect (log-odds: +0.024 per µM; χ² p = 0.0018). The two stimuli acted independently without significant interaction on healthy ephyra production (χ² p = 0.87), although their interaction on total ephyra production (including unhealthy releases) was significant (χ² p = 1.0 × 10^−9^). Notably, low indomethacin concentrations (10 and 25 µM) significantly increased the proportion of unhealthy ephyrae relative to seawater controls (paired t-tests: p = 0.035 and p = 0.006, respectively), while higher concentrations (50–75 µM) did not. Polyp survivorship declined significantly at 28°C and 31°C but was unaffected by indomethacin. Trial replicate was the most statistically significant predictor of every outcome measured with the third trial producing near-universal strobilation but zero viable ephyrae, a result of an unidentified perturbation. These results indicate that temperature elevation to 28–31°C is the most reliable induction strategy for *C. xamachana* and that low indomethacin concentrations should be avoided when ephyra quality is paramount. We also note that cryptic batch-level variables can overwhelm controlled factors and should be addressed in future experimental designs.

## INTRODUCTION

True jellyfish (class Scyphozoa) exhibit a biphasic life cycle comprising a sessile benthic polyp stage and a free-swimming medusa stage. The transition between these stages, strobilation, involves a dramatic metamorphic reorganization in which the polyp’s oral disc differentiates into one or more future medusae (ephyrae) that are sequentially released into the water column(Wang et al. 2020). In monostrobilate species such as *Cassiopea*, only a single ephyra is produced per strobilation event, making the life stage transition essentially binary and tractable to study at the level of individual polyps (Hofmann, Fitt, and Fleck 1996). Strobilation is consequential at multiple scales: in nature, the timing and intensity of polyp strobilation governs the magnitude and phenology of jellyfish blooms, which are intensifying globally in association with coastal eutrophication, ocean warming, and the removal of predators (Schäfer et al. 2021). In the laboratory, reproducible ephyra production is critical for experimental work on cnidarian physiology, symbiosis, and development. Despite its importance, the variables governing strobilation remain incompletely understood, with documented roles for temperature, salinity, light, and multiple chemical cues in different species(Schäfer et al. 2021; Rahat and Adar 1980). The Upside-Down Jellyfish, *Cassiopea xamachana*, has emerged as an important cnidarian model system, particularly for studies of the cnidarian-dinoflagellate symbiosis and for developmental biology (Ohdera et al. 2018). Its tolerance of captive conditions, rapid polyp proliferation through asexual budding, and monostrobilate ephyra production make it logistically convenient for laboratory experiments (Ohdera et al. 2018). Ephyrae of *C. xamachana* are routinely induced in culture using indole compounds, most notably indomethacin (1-(4- chlorobenzoyl)-5-methoxy-2-methyl-1H-indole-3-acetic acid), a non-steroidal anti-inflammatory drug that reliably triggers strobilation across a broad range of scyphozoan taxa (Kuniyoshi et al. 2012; Cabrales-Arellano et al. 2017; Helm and Dunn 2017; Yamamori et al. 2017; Wang et al. 2020; Deng et al. 2022). Temperature shock is also widely used, either alone or in combination with chemical induction, but the relative efficacy of these two approaches and their interaction on ephyra yield and quality has not been systematically quantified in a factorial framework for *Cassiopea*.

A practical concern motivated this work; indomethacin-induced strobilation can produce malformed or non-viable ephyrae; Deng et al. (2022) documented developmental abnormalities in aposymbiotic *C. andromeda* ephyrae induced with indole compounds, and Mostovshchikova et al. ( 2022) reported radial symmetry defects and stunted growth in chemically induced *Aurelia aurita* ephyrae. Whether the concentration of indomethacin modulates the frequency or severity of such abnormalities in *C. xamachana*, and whether temperature provides an alternative induction route with fewer adverse outcomes, is unknown. This is of practical relevance: researchers using chemically induced ephyrae as experimental subjects need to know whether their induction protocol systematically biases the phenotype of the animals they study.

Here we report a full factorial experiment examining five indomethacin levels (0, 10, 25, 50, and 75 µM) × four temperatures (22, 25, 28, and 31°C) in three replicate trials. We quantify the effects of both stimuli, independently and in combination, on (1) ephyra release rate, (2) ephyra viability (growth rate over 14 days post-release), (3) polyp survivorship, (4) day of first ephyra release, and (5) strobilation duration. We additionally characterize a striking trial-level batch effect that, in one trial, overrode treatment-level signals entirely and produced near-universal strobilation with zero viable ephyrae, raising questions about reproducibility that will be relevant to any laboratory maintaining *Cassiopea* cultures.

## METHODS

### Experimental animals and culture

Lab-reared *Cassiopea xamachana* polyps from a non-clonal Floridian founder stock (representative voucher sequence: NCBI Accession No. MZ343250) were maintained in 35 ppt Instant Ocean artificial seawater (ASW) at 22°C on a 12:12 h light:dark cycle at approximately 60 µmol photons m^−2^ s^−1^. Polyps were fed four 24 h-old *Artemia* nauplii per individual per day. Prior to the experiment, polyps were maintained for at least two weeks under these conditions to confirm health and symbiont presence.

### Experimental design

We employed a crossed factorial design with five indomethacin levels (0, 10, 25, 50, and 75 µM) and four temperatures (22, 25, 28, and 31°C). The two zero-indomethacin groups served as controls: an ASW-only group (no solvent) and a DMSO vehicle control group (0.01% DMSO, carrier solvent for indomethacin). Each treatment combination was replicated across three sequential experimental trials (Trial 1: October–December 2022; Trial 2: January–March 2023; Trial 3: March–April 2023), for a cumulative total of 288 individual polyps (96 per trial).

Within each trial, polyps were randomly assigned to individual wells of 24-well plates (one polyp per well, 3 mL of treatment solution per well). Treatment solutions were prepared fresh by dissolving indomethacin (Sigma-Aldrich, ≥99%) in DMSO to a stock concentration of 100 mM, then diluting into ASW to achieve final experimental concentrations. Allocation of polyps to treatment wells was randomized. Group sizes followed an unbalanced design: 9 polyps per group for 0 µM (ASW), 0 µM (DMSO), and 75 µM groups at each temperature; and 15 polyps per group for 10, 25, and 50 µM groups at each temperature (see Table S1 for full allocation).

Four temperature-controlled water baths were used to maintain target temperatures. To account for potential water bath effects as a confounding variable, the temperature assignment of each water bath was rotated between trials (Trials 1 and 2: baths assigned in order 22, 25, 28, 31°C; Trial 3: baths rotated 25, 22, 28, 31°C). Recorded daily mean temperatures and observed ranges for each bath and trial are reported in Table 1. Well plates were held in the baths at all times except during feeding and water changes, which were performed daily (50% water changes) and weekly (100% water changes with cleaning of wells). Total time out of baths was logged daily and averaged < 55 min per day across all trials.

**Table 1.** Recorded daily mean temperatures (°C) and overall range for each target temperature setting across the three experimental trials.

| Target temperature | Trial 1 mean °C (range) | Trial 2 mean °C (range) | Trial 3 mean °C (range) |
| --- | --- | --- | --- |
| 22°C | 22.7 (21.3–23.7) | 21.5 (20.8–22.1) | 21.9 (21.4–22.7) |
| 25°C | 24.9 (24.0–25.1) | 24.8 (24.6–25.1) | 24.2 (22.4–26.3) |
| 28°C | 28.1 (27.7–28.4) | 28.2 (26.7–29.7) | 27.7 (26.6–28.5) |
| 31°C | 31.4 (30.5–31.8) | 30.8 (29.7–31.8) | 30.5 (28.2–31.6) |

### Observation and measurement

All polyps were photographed daily on a Leica MDG41 stereo microscope fitted with a Leica MC170 HD camera (Leica Microsystems, Wetzlar, Germany), yielding approximately 8,900 images across the three trials. Images were analyzed in ImageJ v1.53e (NIH) to measure calyx diameter (polyp) at the widest point. Strobilation was staged daily using the seven-point scheme of Cabrales-Arellano et al. (2017): Stage 1 = resting (no visible strobilation); Stage 2 = calyx widening and flattening; Stage 3 = initial constriction; Stage 4 = deepened constriction; Stage 5 = ephyra semi-formed; Stage 6 = ephyra unfurling; Stage 7 = ephyra complete and pulsing prior to release (see Fig. 1). The day of first strobilation onset was defined as the first day of Stage 2 observation, and strobilation duration was the interval from Stage 2 to the day of ephyra release. The bud-strobilation gap was the number of days between the last recorded asexual bud and the first day of Stage 2.

**Fig. 1.**
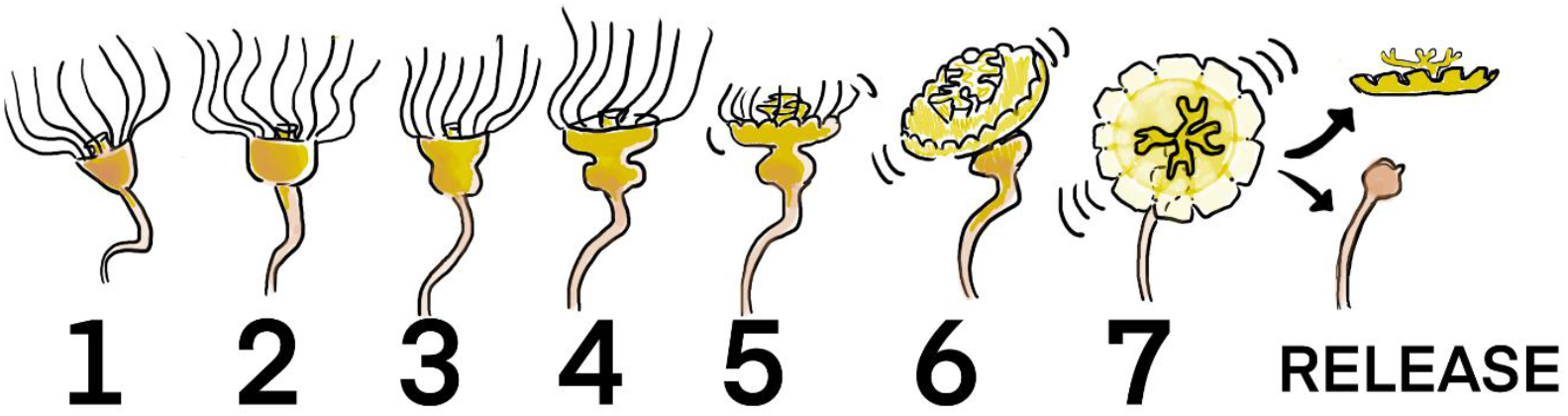
Stages of *Cassiopea* ephyra release as treated in this paper. Stage 1 = resting (no visible strobilation); Stage 2 = calyx widening and flattening; Stage 3 = initial constriction; Stage 4 = deepened constriction; Stage 5 = ephyra semi-formed; Stage 6 = ephyra unfurling; Stage 7 = ephyra complete and pulsing prior to release.

Polyp mortality was defined as complete disintegration of the calyx. Polyps that produced a second ephyra were noted but secondary releases were excluded from all downstream analyses, as secondary ephyrae were almost universally malformed.

At time of daily water exchange, all buds produced by each polyp were counted and removed.

### Ephyra viability assay

Detached ephyrae were immediately transferred to individual wells containing 35 ppt ASW without indomethacin and maintained at room temperature (23°C) on a 12:12 h light:dark cycle. Each ephyra was fed eight *Artemia* nauplii per day. Bell diameter at release was measured from the day-of-release photograph. Subsequent photographs were taken on days 1, 2, 3, 7, and 14 post-release. Ephyra viability (health) was defined as a minimum 30% increase in bell diameter by day 14 relative to release size, a threshold validated against natural growth trajectories reported by Muffett et al. ( 2022). Ephyrae that died, shrank, or grew less than 30% were classified as unhealthy.

### Statistical analyses

Logistic regression models were fitted in R v4.2 to test the effects of indomethacin concentration, temperature, trial, and their pairwise interactions on (i) the probability of any ephyra release and (ii) the probability of a healthy ephyra release. Model fit was evaluated using likelihood ratio tests (χ² statistic). Pairwise comparisons of indomethacin groups to the ASW-only control for unhealthy ephyra production and polyp survival were performed using paired two-sided t-tests (α = 0.05). Each term in the logistic models was assessed by a likelihood-ratio test adjusted for the other terms in the model with trial fitted as a covariate. Correlation between asexual bud count and day of strobilation onset was assessed by Pearson’s correlation. Data visualization used the *ggplot2*, *ggbeeswarm*, and *ggpubr* packages. All analysis code and raw data tables are provided in Supplementary Files S1 (source data) and S2 (analysis code).

## RESULTS

### Overview of ephyra and polyp health outcomes

Across all 288 polyps and three trials, 207 (71.9%) released at least one ephyra within the 29-day experimental window. Of these releases, 96 (33.3% of all polyps; 46.4% of releasing polyps) were classified as healthy and 108 (37.5% of all polyps) were classified as unhealthy. Three additional polyps produced more than one ephyra; these secondary releases were excluded from all analyses. One hundred and nine polyps (37.8%) survived to day 29. Summary statistics stratified by trial, temperature, and indomethacin treatment are presented in Table 2.

**Table 2.** Summary outcomes by trial, temperature, and indomethacin concentration. Values in parentheses are percentages of group total. Ephyra growth ratio = bell diameter at day 14 / bell diameter at release. Dashes (—) indicate statistics not computed for the pooled group due to confounding across trials. * Trial 3 was excluded from most inferential analyses (see text).

| Treatment | n | Any release (%) | Healthy release (%) | Unhealthy release (%) | Polyp survived day 29 (%) | Mean day of release | Mean ephyra growth ratio |
| --- | --- | --- | --- | --- | --- | --- | --- |
| Trial 1 | 96 | 43 (44.8) | 23 (24.0) | 17 (17.7) | 53 (55.2) | 17.9 | 1.26 |
| Trial 2 | 96 | 81 (84.4) | 73 (76.0) | 8 (8.3) | 36 (37.5) | 13.8 | 1.49 |
| Trial 3* | 96 | 83 (86.5) | 0 (0.0) | 83 (86.5) | 20 (20.8) | 11.8 | 0.48 |
| — By temperature — |  |  |  |  |  |  |  |
| 22°C | 72 | 28 (38.9) | 11 (15.3) | 17 (23.6) | 47 (65.3) | — | 0.70 |
| 25°C | 72 | 54 (75.0) | 23 (31.9) | 30 (41.7) | 37 (51.4) | — | 1.04 |
| 28°C | 72 | 64 (88.9) | 30 (41.7) | 32 (44.4) | 14 (19.4) | — | 1.10 |
| 31°C | 72 | 61 (84.7) | 32 (44.4) | 29 (40.3) | 11 (15.3) | — | 1.15 |
| — By indomethacin — |  |  |  |  |  |  |  |
| 0 µM (ASW/DMSO) | 72 | 35 (48.6) | 19 (26.4) | 16 (22.2) | 54 (75.0) | — | — |
| 10 µM | 60 | 43 (71.7) | 19 (31.7) | 24 (40.0) | 21 (35.0) | — | — |
| 25 µM | 60 | 50 (83.3) | 18 (30.0) | 31 (51.7) | 14 (23.3) | — | — |
| 50 µM | 60 | 49 (81.7) | 25 (41.7) | 22 (36.7) | 14 (23.3) | — | — |
| 75 µM | 36 | 30 (83.3) | 15 (41.7) | 15 (41.7) | 6 (16.7) | — | — |

### Dominant effect of trial on all outcome variables

Trial identity was the single strongest predictor of every measured outcome (χ² p < 2.2 × 10^−16^). The three trials differed dramatically in ephyra yield and quality (Table 2; Fig. 2). Trial 1 (autumn 2022) had the lowest strobilation rate (44.8% of polyps released) but the best polyp survivorship (55.2%). Trial 2 (early spring 2023) achieved an 84.4% release rate with 76.0% healthy ephyrae and intermediate survivorship (37.5%). Trial 3 (late spring 2023) produced the highest strobilation rate (86.5%) but with 100% of released ephyrae classified as unhealthy: all failed to grow, most failed to pulse, and many showed signs of arrested development. Polyp mortality in Trial 3 was also substantially higher (20.8% survived to day 29) than in Trial 1.

**Fig. 2.**
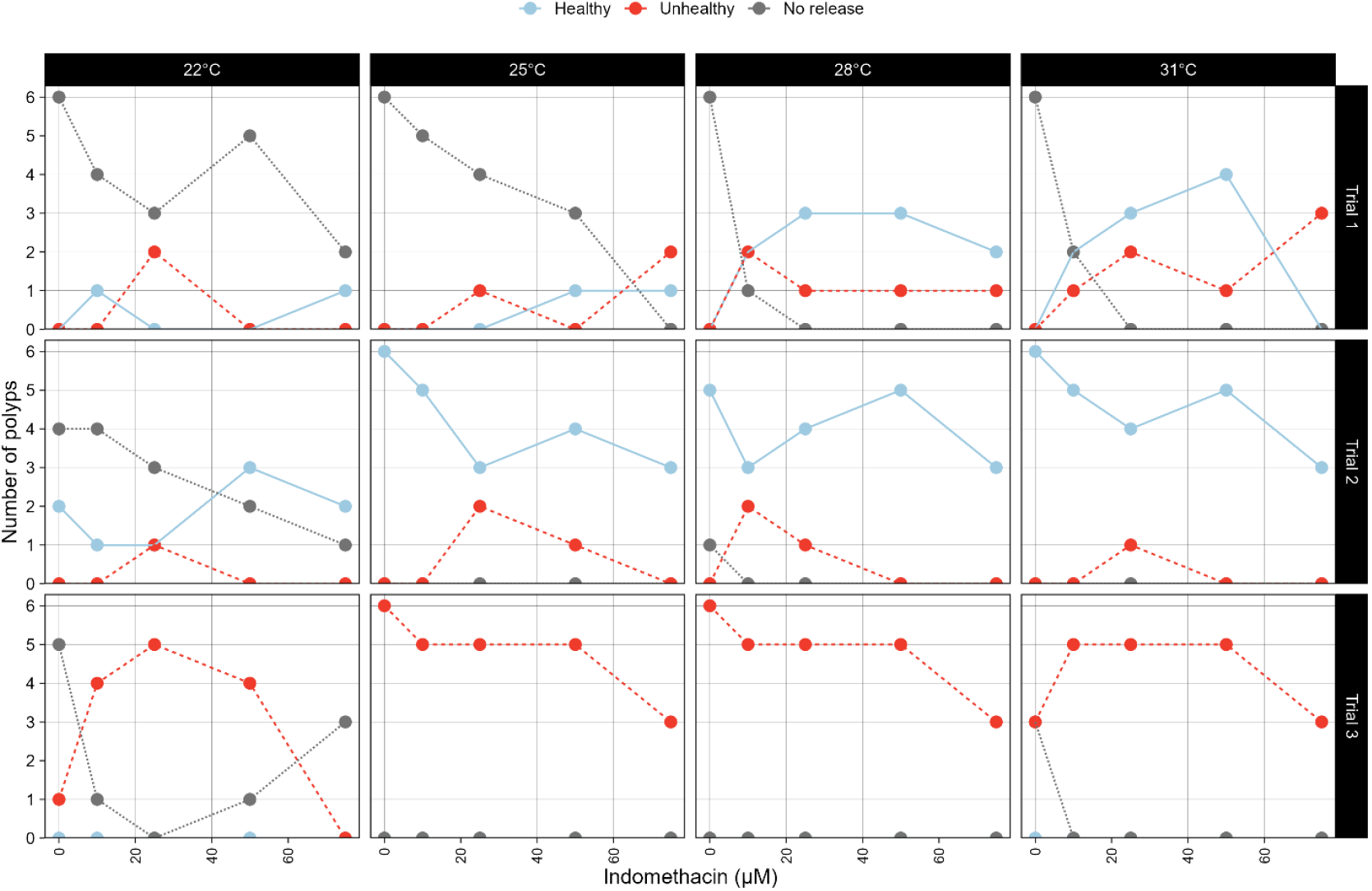
Polyps producing healthy (blue), unhealthy (red) or no (gray) ephyrae by indomethacin treatment. Outcomes are subdivided by treatment temperature (horizontal facet) and trial (vertical facet).

Mean day of ephyra release shortened progressively across trials: Trial 1, 17.9 days (95% CI: 16.4–19.5); Trial 2, 13.8 days (95% CI: 12.5–15.1); Trial 3, 11.8 days (95% CI: 10.9–12.6).

The characteristics of Trial 3, near-universal strobilation accompanied by complete ephyra non-viability and elevated polyp mortality were not explained by any measured parameter, as such, Trial 3 is described separately in the Discussion.

### Effects of temperature on strobilation and ephyra health

Temperature was the dominant controlled predictor of strobilation success. The log-odds of releasing a healthy ephyra increased significantly with temperature (log-odds: +0.327 per °C; χ² = 31.6, df = 1, *p* = 1.9 × 10^−8^). Release rates doubled from 38.9% at 22°C to 88.9% at 28°C, and the proportion of healthy releases increased from 15.3% at 22°C to 41.7–44.4% at 28–31°C (Table 2; Fig. 3). Ephyra growth ratio also improved with temperature: the mean ephyra produced by 22°C polyps shrank after release (14d/release size ratio 0.70), unlike higher temperatures (1.04, 1.10, and 1.15 at 25, 28, and 31°C respectively). Ephyrae from 31°C groups grew significantly faster than those from 22°C groups (paired t-test on trial means, t = 5.83, df = 2, *p* = 0.028).

**Fig. 3.**
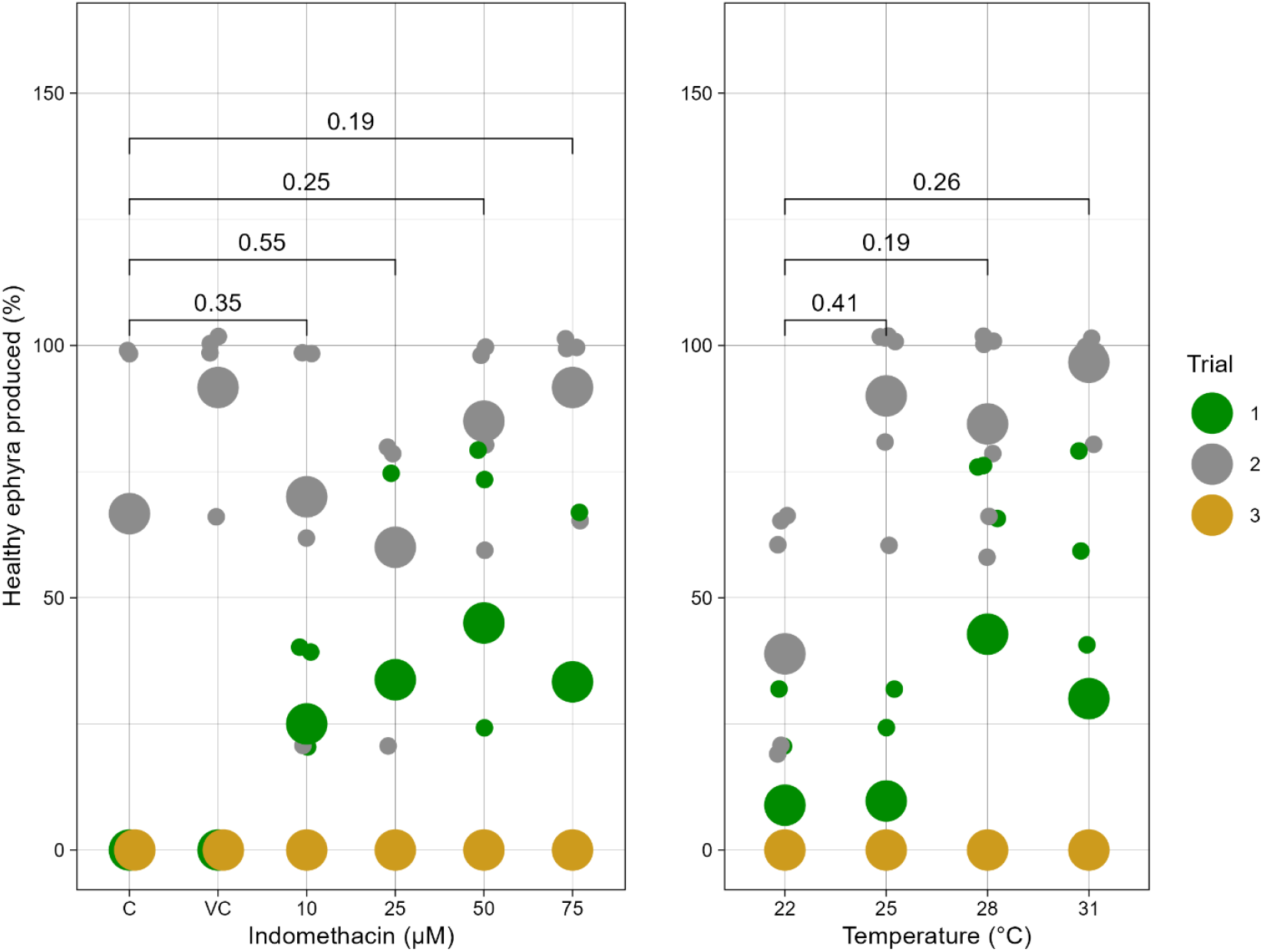
Proportion of polyps releasing a healthy ephyra across all three trials, broken down by indomethacin concentration (left panel) and temperature (right panel); the y-axis of both panels is the percentage of polyps releasing a healthy ephyra. Points represent trial means, jittered for visibility.

Temperature also significantly reduced polyp survivorship at 28°C and 31°C. In Trials 1 and 2 combined, survivorship to day 29 declined from 83.3% of polyps at 22°C to 29.2% at 28°C and 22.9% at 31°C (paired t-tests on trial means, computed across all three trials, relative to 22°C: 28°C, *p* = 0.013; 31°C, *p* = 0.037). This reflects the intrinsic cost of strobilation to polyp integrity: most polyps that released ephyrae died in the weeks following. Polyps in high-temperature groups occasionally survived strobilation and began re-feeding, but commonly went on to release a second, invariably malformed, ephyra.

Day of release was shortened at higher temperatures (Fig. 4), though differences between the elevated temperature groups did not reach statistical significance after accounting for trial. At 22°C, many polyps in ASW/DMSO controls did not strobilate within 29 days in Trial 1, severely limiting comparison of indomethacin effects on timing at that temperature.

**Fig. 4.**
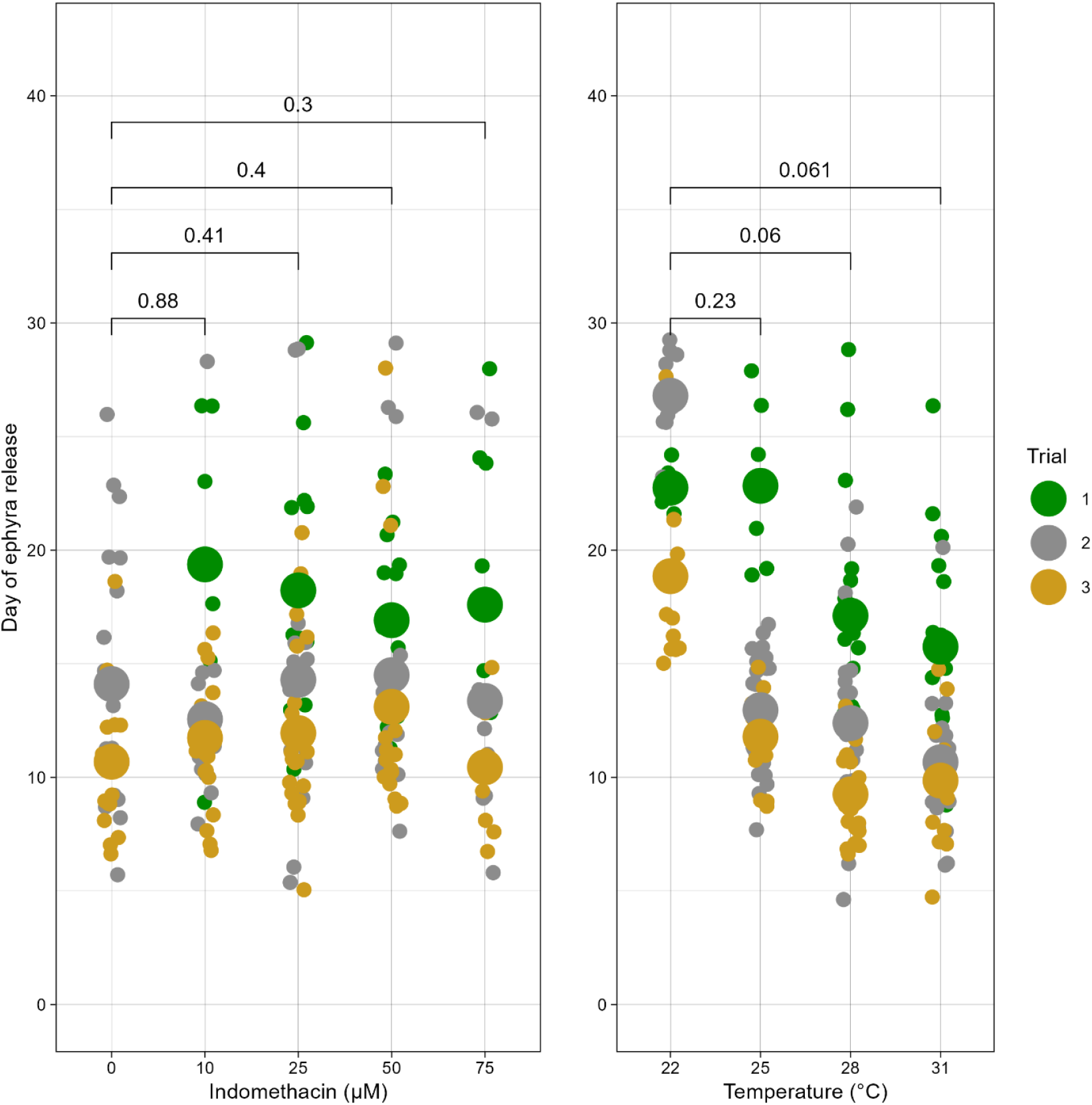
Day of first ephyra release for individual polyps, grouped by indomethacin (left panel) and temperature (right panel), with all trials combined. Points are individual polyps; filled circles are group means. Note that no polyps in 0 µM groups released ephyrae in Trial 1 at any temperature, contributing to the apparent indomethacin-independent release timing in Trials 2 and 3.

### Effects of indomethacin on strobilation and ephyra health

Indomethacin significantly increased the probability of any ephyra release as part of its interaction with temperature (χ² = 37.3, df = 1, *p* = 1.0 × 10^−9^). Even at the lowest concentration tested (10 µM), release rate increased from 48.6% in the zero-indomethacin control to 71.7%, and at 25–75 µM the release rate plateaued around 81–83% (Table 2).

For healthy ephyra production specifically, the effect of indomethacin was smaller and non-monotonic. The log-odds of a healthy release increased with indomethacin concentration (log- odds: +0.024 per µM; χ² = 9.7, df = 1, *p* = 0.0018), but this was driven primarily by the jump in healthy release rate at 50–75 µM (41.7% healthy in both groups) relative to lower concentrations (Table 2). Indomethacin and temperature did not interact significantly in their effect on healthy ephyra probability (df = 1, χ² = 0.025, *p* = 0.874), indicating that they act additively on this outcome.

Critically, low indomethacin concentrations (10 and 25 µM) produced significantly higher proportions of unhealthy ephyrae than the ASW-only control (paired t-tests on trial means: 10 µM, *p* = 0.035; 25 µM, *p* = 0.006; Fig. 5). This pattern was not observed for 50 or 75 µM. The proportion of unhealthy ephyrae was thus non-linearly related to indomethacin concentration, being elevated at intermediate-low concentrations. Indomethacin amount alone was not a significant predictor of the likelihood of an unhealthy (vs. healthy) ephyra among those released (χ², *p* > 0.05), suggesting the elevated unhealthy proportion at 10–25 µM results mainly from increased total release of ephyrae in these groups, including releases of ephyrae that would not otherwise have been produced.

**Fig. 5.**
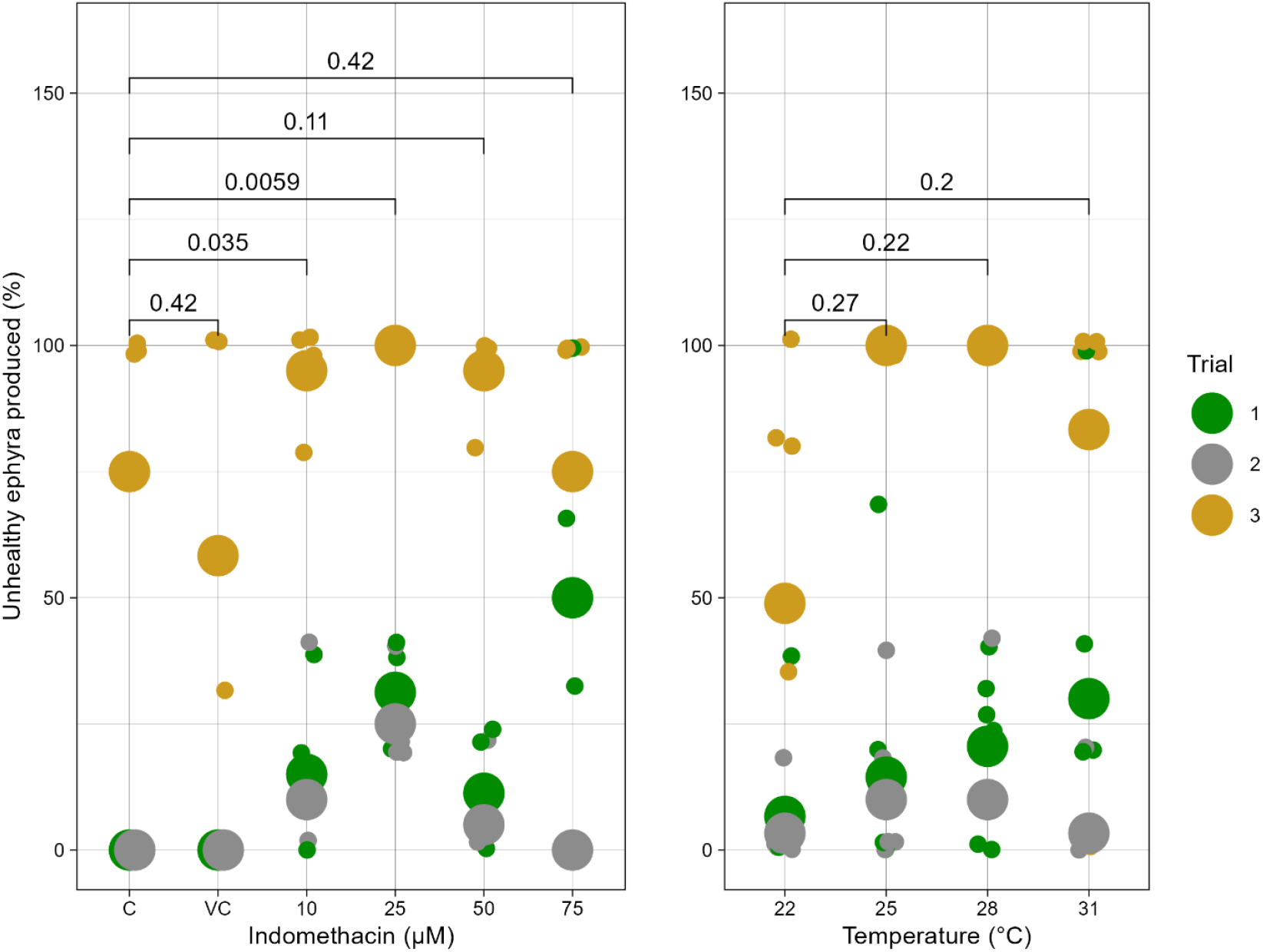
Proportion of polyps releasing an unhealthy ephyra by indomethacin concentration and temperature group, colored by trial. Large central circles indicate group means per trial, small circles indicate treatment-trial means. Horizontal bars indicate significant pairwise differences from the control group (paired t-tests: 10 µM, *p = 0.035; 25 µM,* * p = 0.006).

Indomethacin had no significant effect on polyp survivorship to day 29 (paired t-tests: all p > 0.05 vs. control), though a non-significant downward trend was observed with increasing concentration (survivorship ranging from 75.0% at 0 µM to 16.7% at 75 µM; Table 2, Fig. 6). This trend is partially explained by the greater strobilation rate in indomethacin-treated polyps, since strobilation itself can be lethal or damaging to the polyp.

**Fig. 6.**
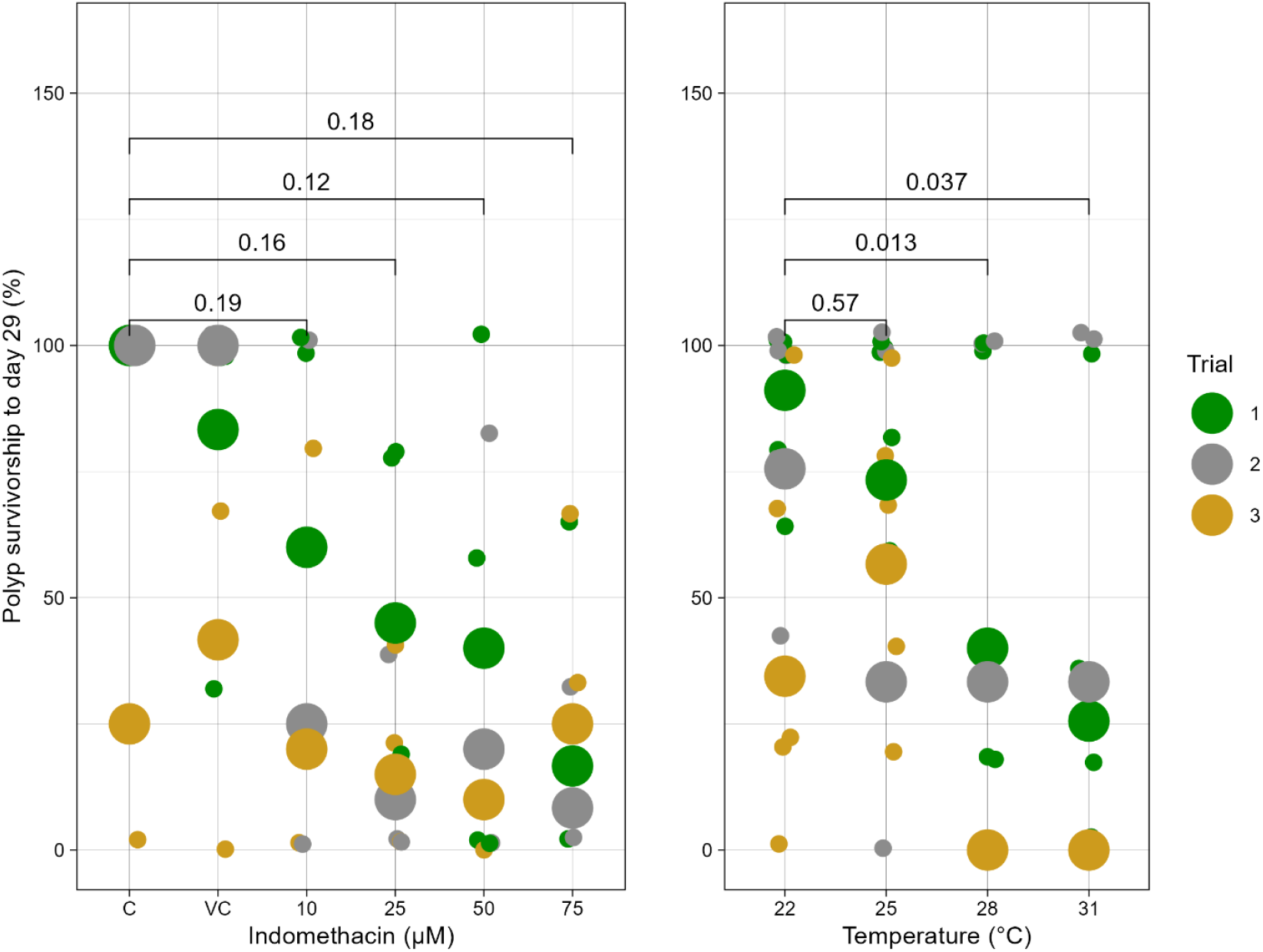
Polyp survivorship to day 29 by indomethacin concentration and temperature group, colored by trial. Large central circles represent group means while small circles represent treatment means within each trial. Horizontal bars indicate significant differences from the 22°C group (paired t-tests: 28°C, *p = 0.013; 31°C,* p = 0.037) or 0 µM (none significant).

### Strobilation timing and bud production

Across all polyps that strobilated (n = 153), strobilation duration (Stage 2 onset to ephyra release) averaged 4.5 days (range: 1–13 days). The bud-strobilation gap (time from last recorded asexual bud to Stage 2 onset) averaged 10.2 days (median: 8 days; interquartile range: 6–13 days). Both of these metrics were highly variable among individuals and did not show systematic relationships with treatment in this dataset.

Total asexual bud production (normalized to starting calyx diameter) showed a non-significant negative trend with both increasing temperature and indomethacin. This pattern is explained by a positive correlation between normalized bud count and the day of strobilation onset (Pearson’s r = 0.51, df = 165, *p* = 1.9 × 10^−12^, Fig. 7). Polyps that strobilated earlier had less time to bud, so the reduction in bud counts at high temperatures and indomethacin concentrations reflects the acceleration of strobilation onset rather than a direct suppression of budding.

**Fig. 7.**
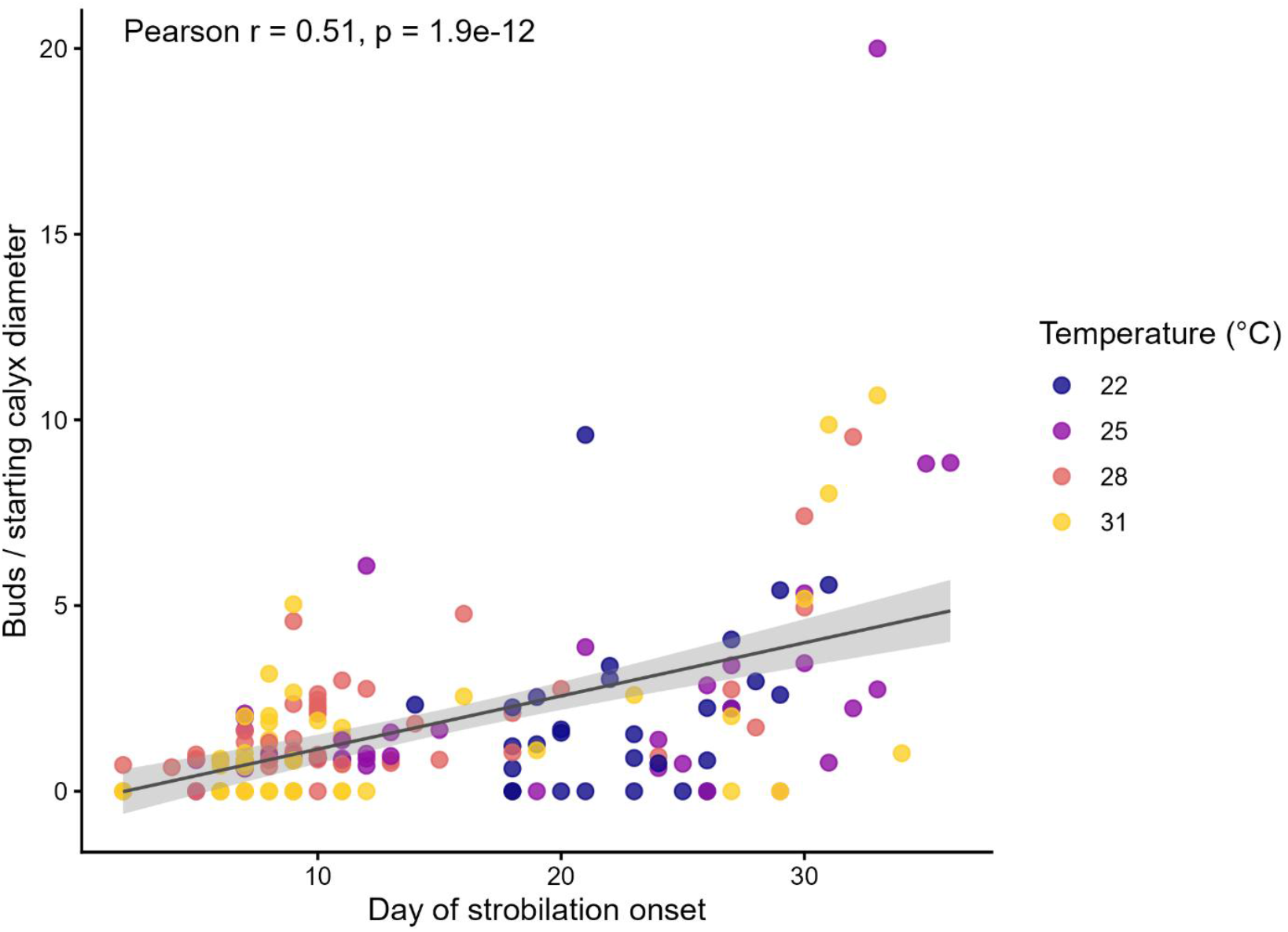
Relationship between number of asexual buds produced (normalized to starting calyx diameter) and the day of strobilation onset for all polyps with complete records (n = 167). Pearson’s r = 0.51, *p* = 1.9 × 10^−12^. Color denotes temperature group. Points are individual polyps; filled circles are group means.

### Size of ephyrae

Ephyra growth was higher in ephyrae originating from 31°C than from 22°C (paired t-test, p = 0.028) (Fig. 8). Ephyra growth was higher with increasing temperature of polyp, despite all ephyrae remaining in ambient temperatures. For trial three, all mean ephyrae growth sat below 1, ephyrae shrank from their size at release. Indomethacin amount had no apparent impact on the health of resultant ephyrae.

**Fig. 8.**
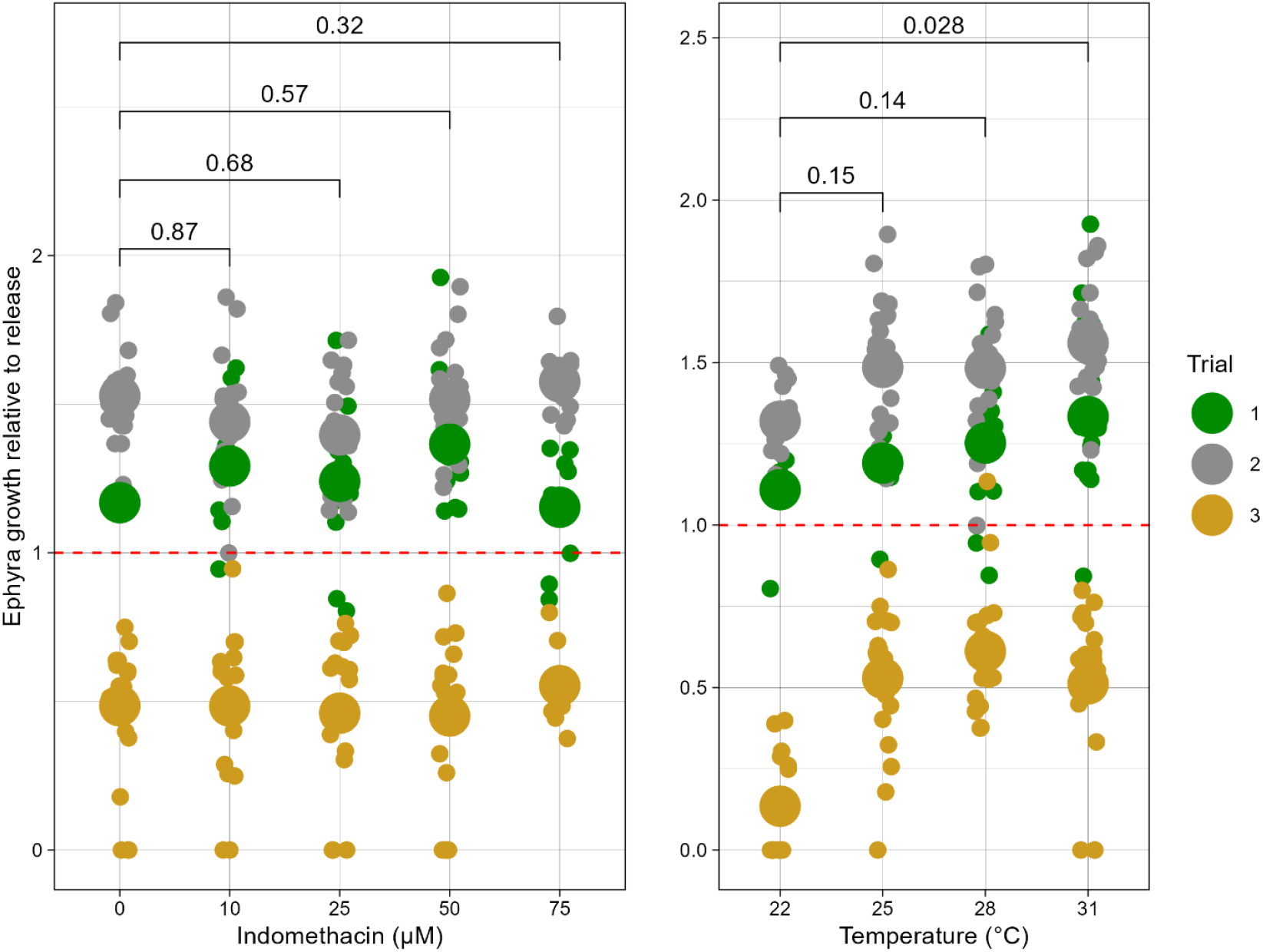
Ephyra growth ratio (bell diameter at day 14 / bell diameter at release) by indomethacin concentration (left) and temperature (right) for all three trials. Ephyrae from Trial 3 shrank (all growth ratio < 1.0). Growth ratio at 31°C was significantly higher than at 22°C (paired t-test, p = 0.028). Points are individual ephyrae; filled circles are group means. Dashed red line represents no growth.

## DISCUSSION

### Temperature outperforms indomethacin as a primary strobilation inducer

Our results establish that temperature elevation is a more reliable and higher-quality driver of strobilation in *Cassiopea xamachana* than indomethacin addition. Raising temperature from the stock-holding condition (22°C) to 28 or 31°C more than doubled the release rate and tripled the proportion of healthy ephyrae, without significantly reducing ephyra growth. In contrast, indomethacin at all concentrations produced smaller gains in healthy ephyra yield, and concentrations below 50 µM *elevated* the rate of unhealthy releases. This is consistent with the observation that indole-induced strobilation in scyphozoans can be associated with developmental abnormalities (Deng et al. 2022, Mostovshchikova et al. 2022), and suggests that for *Cassiopea* specifically, chemically induced metamorphosis at low indole concentrations may be imperfect.

The mechanism by which low indomethacin concentrations produce a worse healthy:unhealthy ratio than higher concentrations is not immediately clear. One possibility is a threshold effect: below a concentration sufficient to coordinate the full strobilation program (i.e., below approximately 50 µM in our system), indomethacin may initiate but not complete the necessary signaling cascade, resulting in premature or aberrant ephyra formation. Cabrales-Arellano et al. (2017) reported that 50 mM was optimal for *C. xamachana* at 25°C, which agrees with our observation that concentrations ≥ 50 µM produced better-quality ephyrae than lower concentrations. For researchers requiring reliable healthy ephyra production, our data support the use of temperature elevation to 28°C or 31°C as the primary induction approach, with 50 µM indomethacin as the preferred chemical adjunct if chemical induction is necessary.

### Interpretations of the interaction between indomethacin and temperature

The significant interaction between indomethacin and temperature for total ephyra production but not for healthy ephyra production alone reflects a pattern in which the two stimuli are partially redundant at high levels: both individually push polyps toward strobilation, and both are required to push reluctant polyps (those at low temperature or in ASW control) toward releasing ephyrae. The lack of interaction on healthy ephyra production suggests that each stimulus independently improves the quality of the strobilation process in a roughly additive manner on the log-odds scale. This additivity implies that the pathways by which temperature and indomethacin act converge on a shared output (healthy medusa formation) through independent upstream routes.

### The batch effect and implications for reproducibility

The dominant trial effect we report, which explained more variance in every outcome variable than temperature or indomethacin, is the most significant finding for practical use of *Cassiopea* as a lab model. The most dramatic manifestation was Trial 3, in which 86.5% of polyps strobilated within a compressed timeframe but produced zero viable ephyrae. Accelerated strobilation with 100% ephyra lethality is, to our knowledge, not previously described in the *Cassiopea* literature and strongly suggests an extreme perturbation. We suspect that, while ’bad batches’ have not been previously reported, they likely occur occasionally for many researchers. As *Cassiopea* polyps are known to associate with specific bacterial communities that vary with symbiotic state (Voolstra and Ziegler 2020), and experimental perturbations of the microbiome can profoundly alter cnidarian physiology, not reporting on a ’bad batch’ that directly impacted our tested physiological outcome would be in poor form. Seasonal variation in water quality, shifts in the bacterial community of aging stock cultures, or the introduction of a transient pathogen could all produce batch-level variation of the magnitude we observed. A similar, milder batch effect differentiated Trial 1 from Trial 2; Trial 1 showing lower release rates and slower strobilation times without producing the catastrophic quality failure seen in Trial 3.

These results have important implications for the use of strobilation data as a quantitative phenotype. If trial-to-trial variation of this magnitude occurs without obvious husbandry differences, then reported strobilation efficiencies from different labs, or even the same lab at different times, may not be directly comparable. We suggest that any strobilation experiment include concurrent negative controls (ASW-only polyps) whose release rates can serve as a retrospective diagnostic: if controls are releasing at high rates, a batch effect is likely. More fundamentally, we advocate for the use of sterile or autoclaved artificial seawater in experimental wells, and for thorough documentation of stock culture history (age, source, recent health events) alongside quantitative strobilation data in published methods. Future work characterizing the microbial communities of polyps during successful versus failed strobilation events would directly test the hypothesis that transient microbiota are a dominant driver of batch variation.

### Practical recommendations

Drawing together our results, we offer the following protocol recommendations for researchers seeking to produce *C. xamachana* ephyrae for experimental use:

1. To maximize the proportion of **healthy ephyrae** with the shortest induction time: **elevate temperature to 28°C** or 31°C without indomethacin addition. This approach yielded healthy release rates of 41.7–44.4% of individual polyps exposed and produced ephyrae with the highest growth rates.
2. To maximize **total ephyra yield** (including some unhealthy ephyrae): combine elevated temperature **(28–31°C) with 50 µM indomethacin**. Release rates in this combination exceeded 85% in Trials 1 and 2.
3. **Avoid indomethacin concentrations of 10–25 µM**: these concentrations increased total release rates but elevated the proportion of unhealthy ephyrae relative to controls, making them suboptimal for most applications.
4. Closely monitor stock culture health and use sterile or UV-treated seawater in experimental wells to minimize the risk of batch effects. If release rates in control wells approach those of treated wells, the experimental batch should be treated with caution.

### Conclusions

We present the first systematic factorial characterization of indomethacin and temperature as co-inducers of strobilation in *Cassiopea xamachana*. Temperature elevation is the more powerful and quality-preserving induction stimulus, while low indomethacin concentrations carry a previously undocumented risk of increasing unhealthy ephyra release. Both stimuli act independently (additively on the log-odds scale) on healthy ephyra production. Despite these clear treatment effects, experimental batch identity explained substantially more variance in strobilation outcomes than any controlled factor. Our results caution against comparing strobilation efficiencies across experiments without concurrent controls and motivate a research agenda aimed at identifying the specific biological variables responsible for batch-to-batch variability in jellyfish culture.

## ACKNOWLEDGMENTS

The authors thank the Miglietta lab for support with animal husbandry and experimental logistics, and members of the Marine Biology Department at Texas A&M University at Galveston for helpful discussions. K.M.M. was supported by a graduate fellowship from Texas A&M University at Galveston. This research was partially supported by NSF award DEB- 2153775 to MPM.

## Supplements

**S1.** Full source data sheet including all primary statistics on ephyra output and survival.

**S2.** Final R code for models and figures.

**S3, S4, S5.** Primary raw data tables from trials 1, 2 and 3.

**S6.** Data sheet used for polyp randomisation.

## LITERATURE CITED

1. Cabrales-Arellano, Patricia, Tania Islas-Flores, Patricia E. Thomé, and Marco A. Villanueva. 2017. “Indomethacin Reproducibly Induces Metamorphosis in Cassiopea Xamachana Scyphistomae.” PeerJ 2017 (3): 1–11. 10.7717/peerj.2979.

2. Deng, Liqiu, Shuhong Wang, Ting Wang, Yuxuan Zhao, and Qi Luo. 2022. “Indoles Can Induce Strobilation in Aposymbiotic Cassiopea Andromeda Polyps but Are Associated with Developmental Abnormalities.” Hydrobiologia 849 (15): 3275–85. 10.1007/s10750-022-04883-z.

3. Helm, Rebecca R., and Casey W. Dunn. 2017. “Indoles Induce Metamorphosis in a Broad Diversity of Jellyfish, but Not in a Crown Jelly (Coronatae).” PLoS ONE 12 (12): 1–13. 10.1371/journal.pone.0188601.

4. Hofmann, Dietrich K., William K. Fitt, and Jürgen Fleck. 1996. “Checkpoints in the Life-Cycle of Cassiopea Spp.: Control of Metagenesis and Metamorphosis in a Tropical Jellyfish.” International Journal of Developmental Biology 40 (1): 331–38. 10.1387/ijdb.8735945.

5. Kuniyoshi, Hisato, Izumi Okumura, Rie Kuroda, Natsumi Tsujita, Kenji Arakawa, Jun Shoji, Tamio Saito, and Hiroyuki Osada. 2012. “Indomethacin Induction of Metamorphosis from the Asexual Stage to Sexual Stage in the Moon Jellyfish , Aurelia Aurita.” Biosci. Biotechnol. Biochem. 76 (7): 1397–1400. 10.1271/bbb.120076.

6. Mostovshchikova, P. S., D. M. Saidov, and I. A. Kosevich. 2022. “Morphological Deviations in Ephyrae after Chemical Induction of Strobilation in Aurelia Aurita (Scyphozoa, Cnidaria).” Russian Journal of Developmental Biology 53 (2): 82–98. 10.1134/s1062360422020084.

7. Muffett, Kaden Mc Kenzie, Joleen Aulgur, and >Maria Pia Miglietta. 2022. “Impacts of Light and Food Availability on Early Development of Cassiopea Medusae.” Frontiers in Marine Science 8 (January): 1–11. 10.3389/fmars.2021.783876.

8. Ohdera, Aki H., Michael J. Abrams, Cheryl L. Ames, David M. Baker, Luis P. Suescún-Bolívar, Allen G. Collins, Christopher J. Freeman, et al. 2018. “Upside-down but Headed in the Right Direction: Review of the Highly Versatile Cassiopea Xamachana System.” Frontiers in Ecology and Evolution 6 (APR): 1–15. 10.3389/fevo.2018.00035.

9. Rahat, M, and Orit Adar. 1980. “Effect of Symbiotic Zooxanthellae and Temperature on Budding and Strobilation in Cassiopea Andromeda.” Biological Bulletin, no. 159: 394–401.

10. Schäfer, Susanne, Sonia K.M. Gueroun, Carlos Andrade, and João Canning-Clode. 2021. “Combined Effects of Temperature and Salinity on Polyps and Ephyrae of Aurelia Solida (Cnidaria: Scyphozoa).” Diversity 13 (11). 10.3390/d13110573.

11. Voolstra, Christian R., and Maren Ziegler. 2020. “Adapting with Microbial Help: Microbiome Flexibility Facilitates Rapid Responses to Environmental Change.” BioEssays 42 (7): 1–9. 10.1002/bies.202000004.

12. Wang, Nan, Minxiao Wang, Yantao Wang, and Chaolun Li. 2020. “Inductive Effect of Bioactive Substances on Strobilation of Jellyfish Aurelia Coerulea.” Journal of Oceanology and Limnology 38 (5): 1548–58. 10.1007/s00343-020-0053-2.

13. Yamamori, Luna, Kazuya Okuizumi, Chika Sato, Shuhei Ikeda, and Haruhiko Toyohara. 2017. “Comparison of the Inducing Effect of Indole Compounds on Medusa Formation in Different Classes of Medusozoa.” Zoological Science 34 (3): 173–78. 10.2108/zs160161.

