## Supplementary material for "Combination of indomethacin and temperature produces reliable strobilation in *Cassiopea xamachana* but batch effects create high variability in ephyra outcomes": S2-S6: S2_Final_codebase_Indomethacin.R

```
library(dplyr)
library(tidyr)
library(ggplot2)
library(ggbeeswarm)
library(ggpubr)
library(cowplot)

data_file <- "S1_Primary_Data_Full_Factorial_Workbook.csv"
Tot <- read.csv(data_file, check.names = TRUE)

# data prep
Tot$Trial <- factor(Tot$Trial)
Tot$facIndomethacin.Level <- factor(Tot$Indomethacin.Level)
Tot$facTemperature <- factor(Tot$Temperature)
Tot$adjBud <- Tot$Bud.num / Tot$Polyp.start.size
Tot$Healthbin <- ifelse(Tot$Healthy.ephyra.produced == "y", 1, 0)
Tot$Probin <- ifelse(Tot$Released == "y", 1, 0)
polypreleases2 <- Tot
bud_ephdia_t12 <- subset(Tot, Trial %in% c("1", "2"))

# aggregated group percentages (C = ASW, VC = DMSO control)
FACT <- Tot %>%
  mutate(Indomethacin = ifelse(Indomethacin.Level == 0 & DMSO == "n", "C",
                               ifelse(Indomethacin.Level == 0 & DMSO == "DMSO", "VC", as.character(Indomethacin.Level)))) %>%
  group_by(Temperature, Indomethacin, Trial) %>%
  summarise(ep.production = 100 * mean(Released == "y"),
            pol.survived = 100 * mean(Survival.through.d29 == "y"),
            Healthy.eph  = 100 * mean(Healthy.ephyra.produced == "y", na.rm = TRUE),
            Unhealthy    = 100 * mean(Healthy.unhealthy.ephyra.produced == "u", na.rm = TRUE), .groups = "drop")
FACT$Indomethacin <- factor(FACT$Indomethacin, levels = c("C", "VC", "10", "25", "50", "75"))
FACT$Temperaturef <- factor(FACT$Temperature)
ReplicateAveragesFACTindo <- FACT %>% group_by(Indomethacin, Trial) %>% summarise(across(c(ep.production, pol.survived, Healthy.eph, Unhealthy), mean), .groups = "drop")
ReplicateAveragesFACTtem  <- FACT %>% group_by(Temperaturef, Trial) %>% summarise(across(c(ep.production, pol.survived, Healthy.eph, Unhealthy), mean), .groups = "drop")

palT <- c("1" = "green4", "2" = "grey55", "3" = "goldenrod3")

# Outcome totals
print(c(n = nrow(Tot), released = sum(Tot$Probin),
        healthy = sum(Tot$Healthbin == 1, na.rm = TRUE),
        unhealthy = sum(Tot$Healthy.unhealthy.ephyra.produced == "u", na.rm = TRUE),
        survived = sum(Tot$Survival.through.d29 == "y")))
#        n  released   healthy unhealthy  survived
#      288       207        96       108       109

# Healthy ephyra logistic regression (sequential LR: indomethacin, temperature, trial)
mH <- glm(Healthbin ~ Indomethacin.Level + Temperature + Trial, data = Tot, family = binomial)
print(summary(mH)); print(anova(mH, test = "LRT"))

# Healthy ephyra indomethacin x temperature interaction
print(anova(mH, glm(Healthbin ~ Indomethacin.Level * Temperature + Trial, data = Tot, family = binomial), test = "LRT"))

# Any-release indomethacin x temperature interaction
mR <- glm(Probin ~ Indomethacin.Level + Temperature + Trial, data = Tot, family = binomial)
print(anova(mR, glm(Probin ~ Indomethacin.Level * Temperature + Trial, data = Tot, family = binomial), test = "LRT"))

# Unhealthy ephyra paired t-tests vs ASW control
print(compare_means(Unhealthy ~ Indomethacin, data = ReplicateAveragesFACTindo, method = "t.test", paired = TRUE, ref.group = "C"))

# Polyp survivorship paired t-tests vs 0
print(compare_means(pol.survived ~ Indomethacin, data = ReplicateAveragesFACTindo, method = "t.test", paired = TRUE, ref.group = "C"))

# Polyp survivorship paired t-tests vs 22C
print(compare_means(pol.survived ~ Temperaturef, data = ReplicateAveragesFACTtem, method = "t.test", paired = TRUE, ref.group = "22"))

# Ephyra growth 31C vs 22C paired t-test
rg <- Tot %>% group_by(facTemperature, Trial) %>% summarise(g = mean(Eph.growth, na.rm = TRUE), .groups = "drop")
print(t.test(rg$g[rg$facTemperature == "31"], rg$g[rg$facTemperature == "22"], paired = TRUE))

# Pearson correlation buds vs strobilation onset
print(cor.test(Tot$Strob_Start, Tot$adjBud))

# Strobilation timing descriptives
print(summary(Tot$Strob_time)); print(summary(Tot$Bud_Strob_gap))

# Fig 1 outcome counts by group, trial and temperature
d <- Tot %>%
  mutate(Outcome = recode(Healthy.unhealthy.ephyra.produced, y = "Healthy", u = "Unhealthy", n = "No release", .default = NA_character_),
         TrialN = as.integer(as.character(Trial))) %>%
  filter(!is.na(Outcome)) %>%
  count(Temperature, Indomethacin.Level, TrialN, Outcome, name = "count") %>%
  complete(nesting(Temperature, Indomethacin.Level), TrialN = 1:3,
           Outcome = c("Healthy", "Unhealthy", "No release"), fill = list(count = 0))
d$Trial <- factor(d$TrialN, levels = 1:3, labels = c("Trial 1", "Trial 2", "Trial 3"))
d$Outcome <- factor(d$Outcome, levels = c("Healthy", "Unhealthy", "No release"))
d$Temp <- factor(d$Temperature); levels(d$Temp) <- c("22°C", "25°C", "28°C", "31°C")
pal2 <- c("Healthy" = "#9ecae1", "Unhealthy" = "#ef3b2c", "No release" = "#737373")
fig1 <- ggplot(d, aes(Indomethacin.Level, count, colour = Outcome, group = Outcome, linetype = Outcome)) +
  geom_point(size = 3) + geom_line() +
  scale_linetype_manual(values = c(Healthy = "solid", Unhealthy = "22", "No release" = "11"), guide = "none") +
  scale_colour_manual(values = pal2, name = NULL) +
  scale_y_continuous(breaks = seq(0, 6, 1), expand = expansion(mult = c(0, .05))) +
  labs(x = "Indomethacin (μM)", y = "Number of polyps") +
  theme_linedraw(base_size = 13) +
  theme(panel.grid.minor = element_blank(),
        axis.text.x = element_text(angle = 90, vjust = .5, hjust = 1, size = 9),
        legend.position = "top") + facet_grid(Trial ~ Temp)
ggsave("Fig1_outcome_counts.png", fig1, width = 12, height = 8, dpi = 300)

# Fig 2 healthy ephyra production
ind_h <- ggplot(FACT, aes(Indomethacin, Healthy.eph, color = Trial)) +
  geom_jitter(width = .1, cex = 3) + ylim(0, 160) +
  scale_colour_manual(name = "Trial", values = palT) +
  geom_beeswarm(data = ReplicateAveragesFACTindo, size = 8) +
  stat_compare_means(data = ReplicateAveragesFACTindo, comparisons = list(c("C", "10"), c("C", "25"), c("C", "50"), c("C", "75")), method = "t.test", paired = TRUE) +
  xlab("Indomethacin (μM)") + ylab("Healthy ephyra produced (%)") + theme_linedraw() + theme(legend.position = "none")
tem_h <- ggplot(FACT, aes(Temperaturef, Healthy.eph, color = Trial)) +
  geom_jitter(width = .1, cex = 3) + ylim(0, 160) +
  scale_colour_manual(name = "Trial", values = palT) +
  geom_beeswarm(data = ReplicateAveragesFACTtem, size = 8) +
  stat_compare_means(data = ReplicateAveragesFACTtem, comparisons = list(c("22", "25"), c("22", "28"), c("22", "31")), method = "t.test", paired = TRUE) +
  xlab("Temperature (°C)") + ylab("") + theme_linedraw()
ggsave("Fig2_healthy_ephyra.png", plot_grid(ind_h, tem_h, ncol = 2), width = 8, height = 6, dpi = 300)

# Fig 3 unhealthy ephyra production
indounh <- ggplot(FACT, aes(Indomethacin, Unhealthy, color = Trial)) +
  geom_jitter(width = .1, cex = 3) + ylim(0, 160) +
  scale_color_manual(name = "Trial", values = palT) +
  geom_beeswarm(data = ReplicateAveragesFACTindo, size = 8) +
  stat_compare_means(data = ReplicateAveragesFACTindo, comparisons = list(c("C", "VC"), c("C", "10"), c("C", "25"), c("C", "50"), c("C", "75")), method = "t.test", paired = TRUE) +
  xlab("Indomethacin (μM)") + ylab("Unhealthy ephyra produced (%)") + theme_linedraw() + theme(legend.position = "none")
tempunh <- ggplot(FACT, aes(Temperaturef, Unhealthy, color = Trial)) +
  geom_jitter(width = .1, cex = 3) + ylim(0, 160) +
  scale_colour_manual(name = "Trial", values = palT) +
  geom_beeswarm(data = ReplicateAveragesFACTtem, size = 8) +
  stat_compare_means(data = ReplicateAveragesFACTtem, comparisons = list(c("22", "25"), c("22", "28"), c("22", "31")), method = "t.test", paired = TRUE) +
  xlab("Temperature (°C)") + ylab("") + theme_linedraw()
ggsave("Fig3_unhealthy_ephyra_release.png", plot_grid(indounh, tempunh, ncol = 2), width = 8, height = 6, dpi = 300)

# Fig 4 polyp survivorship
indosurv <- ggplot(FACT, aes(Indomethacin, pol.survived, color = Trial)) +
  geom_jitter(width = .1, cex = 3) + ylim(0, 160) +
  scale_colour_manual(name = "Trial", values = palT) +
  geom_beeswarm(data = ReplicateAveragesFACTindo, size = 8) +
  stat_compare_means(data = ReplicateAveragesFACTindo, comparisons = list(c("C", "10"), c("C", "25"), c("C", "50"), c("C", "75")), method = "t.test", paired = TRUE) +
  xlab("Indomethacin (μM)") + ylab("Polyp survivorship to day 29 (%)") + theme_linedraw() + theme(legend.position = "none")
temsurv <- ggplot(FACT, aes(Temperaturef, pol.survived, color = Trial)) +
  geom_jitter(width = .1, cex = 3) + ylim(0, 160) +
  scale_colour_manual(name = "Trial", values = palT) +
  geom_beeswarm(data = ReplicateAveragesFACTtem, size = 8) +
  stat_compare_means(data = ReplicateAveragesFACTtem, comparisons = list(c("22", "25"), c("22", "28"), c("22", "31")), method = "t.test", paired = TRUE) +
  xlab("Temperature (°C)") + ylab("") + theme_linedraw()
ggsave("Fig4_survivorship.png", plot_grid(indosurv, temsurv, ncol = 2), width = 8, height = 6, dpi = 300)

# Fig 5 day of ephyra release
ReplicateAverages <- subset(polypreleases2, Released != "n") %>% group_by(facIndomethacin.Level, Trial) %>% summarise(DoR = mean(DoR, na.rm = TRUE), .groups = "drop")
indoDor <- ggplot(subset(polypreleases2, Released != "n"), aes(facIndomethacin.Level, DoR, color = factor(Trial))) +
  geom_jitter(width = .1, cex = 3) +
  scale_colour_manual(name = "Trial", values = palT) +
  geom_beeswarm(data = ReplicateAverages, size = 8) + ylim(0, 42) +
  stat_compare_means(data = subset(ReplicateAverages, Trial != "1"), comparisons = list(c("0", "10"), c("0", "25"), c("0", "50"), c("0", "75")), method = "t.test", paired = TRUE) +
  xlab("Indomethacin (μM)") + ylab("Day of ephyra release") + theme_linedraw() + theme(legend.position = "none")
ReplicateAverages2 <- subset(polypreleases2, Released != "n") %>% group_by(facTemperature, Trial) %>% summarise(DoR = mean(DoR, na.rm = TRUE), .groups = "drop")
temDor <- ggplot(subset(polypreleases2, Released != "n"), aes(facTemperature, DoR, color = factor(Trial))) +
  geom_jitter(width = .1, cex = 3) + ylim(0, 42) +
  scale_colour_manual(name = "Trial", values = palT) +
  geom_beeswarm(data = ReplicateAverages2, size = 8) +
  stat_compare_means(data = ReplicateAverages2, comparisons = list(c("22", "25"), c("22", "28"), c("22", "31")), method = "t.test", paired = TRUE) +
  xlab("Temperature (°C)") + ylab("") + theme_linedraw()
ggsave("Fig5_Day_of_Ephyra_Release.png", plot_grid(indoDor, temDor, ncol = 2), width = 8, height = 8, dpi = 300)

# Fig 6 buds vs strobilation onset
fig6 <- ggplot(subset(Tot, !is.na(Strob_Start) & !is.na(adjBud)), aes(Strob_Start, adjBud, color = facTemperature)) +
  geom_point(size = 3) + geom_smooth(method = "lm", se = TRUE, color = "black") +
  scale_colour_viridis_d(name = "Temperature (°C)", option = "C", end = .9) +
  xlab("Day of strobilation onset") + ylab("Buds / starting calyx diameter") + theme_linedraw()
ggsave("Fig6_buds_vs_strobilation.png", fig6, width = 7, height = 5, dpi = 300)

# Fig 7 ephyra growth relative to release
ReplicateAverages8 <- subset(Tot, !is.na(Eph.growth)) %>% group_by(facIndomethacin.Level, Trial) %>% summarise(Eph.growth = mean(Eph.growth, na.rm = TRUE), .groups = "drop")
indoxx <- ggplot(Tot, aes(facIndomethacin.Level, Eph.growth, color = Trial)) +
  geom_jitter(width = .1, cex = 3) +
  scale_colour_manual(name = "Trial", values = palT) + geom_hline(yintercept=c(1), linetype="dashed", color = "red")+
  geom_beeswarm(data = ReplicateAverages8, size = 8) +
  stat_compare_means(data = ReplicateAverages8, comparisons = list(c("0", "10"), c("0", "25"), c("0", "50"), c("0", "75")), method = "t.test", paired = TRUE) +
  xlab("Indomethacin (μM)") + ylab("Ephyra growth relative to release") + theme_linedraw() + theme(legend.position = "none")
ReplicateAverages9 <- subset(Tot, !is.na(Eph.growth)) %>% group_by(facTemperature, Trial) %>% summarise(Eph.growth = mean(Eph.growth, na.rm = TRUE), .groups = "drop")
temxx <- ggplot(Tot, aes(facTemperature, Eph.growth, color = factor(Trial))) +
  geom_jitter(width = .1, cex = 3) + geom_hline(yintercept=c(1), linetype="dashed", color = "red")+
  scale_colour_manual(name = "Trial", values = palT) +
  geom_beeswarm(data = ReplicateAverages9, size = 8) +
  stat_compare_means(data = ReplicateAverages9, comparisons = list(c("22", "25"), c("22", "28"), c("22", "31")), method = "t.test", paired = TRUE) +
  xlab("Temperature (°C)") + ylab("") + theme_linedraw()
ggsave("Fig7_ephyra_release_size.png", plot_grid(indoxx, temxx, ncol = 2), width = 8, height = 6, dpi = 300)

# SuppFig number of buds
ReplicateAverages3 <- Tot %>% group_by(facIndomethacin.Level, Trial) %>% summarise(adjBud = mean(adjBud, na.rm = TRUE), .groups = "drop")
indoRep <- ggplot(Tot, aes(facIndomethacin.Level, adjBud, color = Trial)) +
  geom_jitter(width = .1, cex = 3) +
  scale_colour_manual(name = "Trial", values = palT) +
  geom_beeswarm(data = ReplicateAverages3, size = 8) +
  stat_compare_means(data = ReplicateAverages3, comparisons = list(c("0", "10"), c("0", "25"), c("0", "50"), c("0", "75")), method = "t.test", paired = TRUE) +
  xlab("Indomethacin (μM)") + ylab("Bud number") + theme_linedraw() + theme(legend.position = "none")
ReplicateAverages4 <- Tot %>% group_by(facTemperature, Trial) %>% summarise(adjBud = mean(adjBud, na.rm = TRUE), .groups = "drop")
temRep <- ggplot(Tot, aes(facTemperature, adjBud, color = factor(Trial))) +
  geom_jitter(width = .1, cex = 3) +
  scale_colour_manual(name = "Trial", values = palT) +
  geom_beeswarm(data = ReplicateAverages4, size = 8) +
  stat_compare_means(data = ReplicateAverages4, comparisons = list(c("22", "25"), c("22", "28"), c("22", "31")), method = "t.test", paired = TRUE) +
  xlab("Temperature (°C)") + ylab("") + theme_linedraw()
ggsave("FigSfig_Number_of_buds.png", plot_grid(indoRep, temRep, ncol = 2), width = 8, height = 8, dpi = 300)

# SuppFig bud-strobilation gap
ReplicateAverages5 <- subset(bud_ephdia_t12, !is.na(Bud_Strob_gap)) %>% group_by(facIndomethacin.Level, Trial) %>% summarise(Bud_Strob_gap = mean(Bud_Strob_gap, na.rm = TRUE), .groups = "drop")
indobg <- ggplot(bud_ephdia_t12, aes(facIndomethacin.Level, Bud_Strob_gap, color = Trial)) +
  geom_jitter(width = .1, cex = 3) + ylim(0, 46) +
  scale_colour_manual(name = "Trial", values = palT) +
  geom_beeswarm(data = ReplicateAverages5, size = 8) +
  stat_compare_means(data = ReplicateAverages5, comparisons = list(c("0", "10"), c("0", "25"), c("0", "50"), c("0", "75")), method = "t.test", paired = TRUE) +
  xlab("Indomethacin (μM)") + ylab("Gap between final bud and strobila") + theme_linedraw() + theme(legend.position = "none")
ReplicateAverages6 <- subset(bud_ephdia_t12, !is.na(Bud_Strob_gap)) %>% group_by(facTemperature, Trial) %>% summarise(Bud_Strob_gap = mean(Bud_Strob_gap, na.rm = TRUE), .groups = "drop")
tembg <- ggplot(bud_ephdia_t12, aes(facTemperature, Bud_Strob_gap, color = factor(Trial))) +
  geom_jitter(width = .1, cex = 3) + ylim(0, 46) +
  scale_colour_manual(name = "Trial", values = palT) +
  geom_beeswarm(data = ReplicateAverages6, size = 8) +
  stat_compare_means(data = ReplicateAverages6, comparisons = list(c("22", "25"), c("22", "28"), c("22", "31")), method = "t.test", paired = TRUE) +
  xlab("Temperature (°C)") + ylab("") + theme_linedraw()
ggsave("FigSfig_bud-strob-gap.png", plot_grid(indobg, tembg, ncol = 2), width = 8, height = 5, dpi = 300)
```
