## Supplementary figures and images for "Combination of indomethacin and temperature produces reliable strobilation in *Cassiopea xamachana* but batch effects create high variability in ephyra outcomes"

### S6_Methods_Well_Plate_Photo.jpg

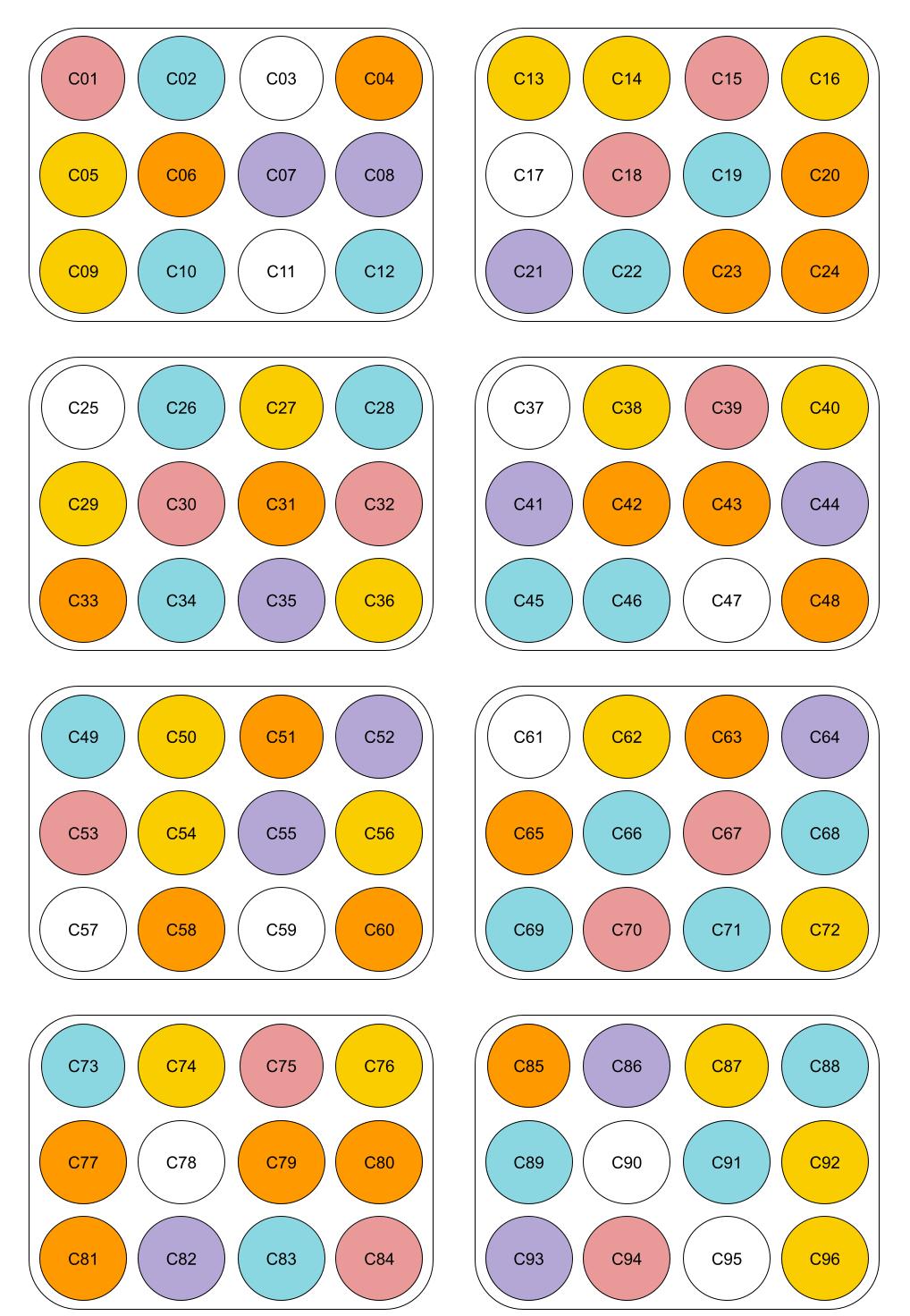
